# Novel CHD8 genomic targets identified in fetal mouse brain by *in vivo* Targeted DamID

**DOI:** 10.1101/2021.01.12.426468

**Authors:** A. Ayanna Wade, Jelle van den Ameele, Seth W. Cheetham, Rebecca Yakob, Andrea H. Brand, Alex S. Nord

**Author notes:** Mater Research Institute-University of Queensland, Woolloongabba, QLD, Australia. These authors contributed equally (A. Ayanna Wade, Jelle van den Ameele). Corresponding authors Correspondence should be directed to: Alex. S. Nord or Andrea H. Brand.

## Abstract

Genetic studies of autism spectrum disorder (ASD) have revealed a causal role for mutations in chromatin remodeling genes. Chromodomain helicase DNA binding protein 8 (*CHD8*) encodes a chromatin remodeler with one of the highest *de novo* mutation rates in sporadic ASD. However, the relationship between CHD8 genomic function and autism-relevant biology remains poorly elucidated. CHD8 binding studies have relied on <u>Ch</u>romatin Immuno<u>p</u>recipitation followed by sequencing (ChIP-seq), however, these datasets exhibit significant variability. ChIP-seq has technical limitations in the context of weak or indirect protein-DNA interactions or when high-performance antibodies are unavailable. Thus, complementary approaches are needed overall, and, specifically, to establish CHD8 genomic targets and regulatory function. Here we used Targeted DamID *in utero* to characterize CHD8 binding in developing embryonic mouse cortex. CHD8 Targeted DamID followed by sequencing (CHD8 TaDa-seq) revealed binding at previously identified targets as well as loci sensitive to *Chd8* haploinsufficiency. CHD8 TaDa-seq highlighted CHD8 binding distal to a subset of genes specific to neurodevelopment and neuronal function. These studies establish TaDa-seq as a useful alternative for mapping protein-DNA interactions *in vivo* and provide insights into the relationship between chromatin remodeling by CHD8 and autism-relevant pathophysiology associated with *CHD8* mutations.

## INTRODUCTION

Neurodevelopmental disorders (NDDs) including autism spectrum disorder (ASD) and intellectual disability (ID) are complex disorders caused by genetic and environmental factors that disrupt brain development. Genetic studies have identified an overlapping set of genes that, when mutated, greatly increase risk for both ASD and ID (1–8). Of these shared risk gene sets, a striking and surprising finding has been the strong enrichment of case mutations in genes that encode proteins involved in chromatin remodeling (1,4). One of these genes, with among the highest number of identified ASD and ID case mutations, is Chromodomain Helicase DNA binding protein 8 (*CHD8*). Characterization of patient phenotypes associated with loss-of-function *CHD8* mutations has revealed a syndrome-like pattern of pathology.

These patients commonly feature symptoms meeting stringent ASD diagnosis, a spectrum of ID and cognitive impairment, macrocephaly, gastrointestinal and sleep disturbances, and other symptoms (9–13). The function of CHD8 and other NDD-associated chromatin remodeling proteins in developing brain remains poorly characterized, representing a major barrier to understanding the neurodevelopmental mechanisms of NDDs.

Chromatin remodelers impact the packaging and functional readout of DNA through interactions with chromatin (14). The dominant approach to understanding molecular function of DNA-associated proteins is to map their specific genomic targets, primarily by <u>Ch</u>romatin <u>I</u>mmuno<u>p</u>recipitation followed by sequencing (ChIP-seq). ChIP-seq has been successfully applied to identify targets of ASD/ID-associated chromatin remodelers, including when profiling fetal brain tissue. However, ChIP-seq requires specific and sensitive antibodies, sufficient sample, and processing steps, specifically crosslinking and fragmentation, that can introduce signal artifacts (15). Further, ChIP-seq performs best with strong, typically direct, interactions between the protein and target DNA (16). This is a significant drawback, as many chromatin remodelers interact indirectly with DNA and ChIP-seq grade antibodies are not always available. Thus, a major limitation to studies of NDD-associated chromatin remodelers has been the challenges presented in identifying genomic interactions by ChIP-seq. One common alternative strategy to ChIP-seq has been to introduce epitope-tagged versions of these proteins to improve immunoprecipitation (17). While this strategy overcomes some barriers, often there are still technical obstacles. For example, epitope tags may address the lack of ChIP-seq grade antibodies, but issues still remain for weak or indirect protein-DNA interactions and artifacts introduced by crosslinking and fragmentation (16).

The growing list of studies that report ChIP-seq derived genomic binding patterns of CHD8 across human and mouse brain tissues and *in vitro* models exemplifies the challenges of applying ChIP-seq to understand chromatin remodeler function (18–24). Our meta-analysis of published CHD8 ChIP-seq datasets found strong concordance across datasets for the strongest genomic interactions (25). However, there was extensive variability in the number and genomic distribution of CHD8 ChIP-seq peaks. This was true even among studies that examined similar tissue types, e.g. adult mouse cortex (21–23), and for studies that used the same antibodies and general methods. Thus, biological inferences regarding CHD8 function have varied considerably based on which ChIP-seq dataset is used. This is reflected in CHD8 publications that highlight various patterns: at one end, widespread binding including at the majority of promoters (19,23,24); at the other end, more limited binding primarily at promoters of genes involved in basic cell functions (21–22). These contrasting ChIP-seq findings demonstrate the need for complementary methods to map genomic interactions for CHD8 and, more generally, for chromatin remodelers and other difficult to ChIP proteins.

Motivated by the need for approaches that avoid antibody-based limitations and technical issues that can be associated with ChIP-seq, we decided to use Targeted DamID (TaDa) (26–27) to map CHD8 targets *in vivo* in fetal mouse cortex. In TaDa, a protein of interest (here CHD8) is fused to an E. coli DNA adenine methyltransferase domain (Dam). Wherever the Dam fusion protein interacts with the genome, the methylase catalyzes methylation of adenine within the sequence GATC. As endogenous adenine methylation is extremely rare in eukaryotes (28–31), the genomic interaction targets of the protein of interest can be identified by mapping adenine methylation in the genome. This approach does not require cell sorting, fixation, crosslinking, or affinity purification, as interactions are mapped via restriction digestion at methylated GATC sites, followed by DNA sequencing (32). TaDa has been used successfully to map genome-wide binding of transcription factors, chromatin proteins and RNA polymerase in *Drosophila* and mammalian cells (for example, 26-27, 33-37) as well as to map non-coding RNA interactions with the genome (38).

Here, we delivered TaDa constructs by *in utero* electroporation (IUE) to perform CHD8 TaDa-seq in the developing mouse brain *in vivo*. Our results show the feasibility and value of this approach, resolving CHD8 interactions in embryonic mouse cerebral cortex. More broadly, our study highlights a novel approach towards mapping genomic binding patterns of proteins that are challenging or intractable to ChIP-seq.

## MATERIALS AND METHODS

### Targeted DamID Constructs

Previously, we developed Targeted DamID (TaDa) to enable cell type-specific profiling *in vivo* while avoiding the potential toxicity resulting from expression of high levels of Dam methylase (26). Using TaDa, transcription of a primary open reading frame (ORF1; here mCherry) is followed by two TAA stop codons and a single nucleotide frameshift upstream of a secondary open reading frame: the coding sequence of the Dam fusion protein (ORF2; here Dam-CHD8). Translation of this bicistronic message results in expression of ORF1 as well as extremely low levels of the Dam fusion protein (ORF2) due to rare ribosomal re-entry and translational re-initiation. TaDa enables rapid, accurate and sensitive identification of genomic binding sites.

When Dam-fusion proteins are expressed in *dam*^*–*^ bacteria, the methylase is able to methylate plasmid DNA. In transient transfection experiments, methylated plasmid DNA co-amplifies with genomic DNA and constitutes a substantial proportion of the sequencing library. For this reason, DamID was thought to be incompatible with transient transfection (39). We introduced an intron into the coding sequence of the Dam methylase to prevent expression in bacteria but not in eukaryotes, where the intron is removed and the enzyme is expressed (J.v.d.A., S.W.C. and A.H.B., unpublished).

To generate the experimental plasmid, pCAG-mCherry-intronDam-CHD8, encoding the Dam methylase fused to the human *CHD8* open reading frame (hereafter CHD8 TaDa), a full-length CHD8 isoform (Origene, RG230753) was subcloned by Gibson assembly into pCAG-mCherry-intronDam, C-terminal to the Dam methylase and a myc-tag. The control plasmid was pCAG-mCherry-intronDam (hereafter Dam-only). Plasmids were sequenced following subcloning. pCAG-Venus, encoding a variant of green fluorescent protein, served as a control for efficiency of *in utero* electroporation and to enable dissection of the electroporated region.

### Delivery via IUE of fetal mouse cortex and generation of TaDa libraries for sequencing

MF1 mice from the same litter were *in utero* electroporated as previously described (40–41). CHD8 TaDa (0.5ug/ul) or Dam-only (0.5ug/ul) and electroporation-control (0.25ug/ul) plasmids were injected into the fetal brain ventricles at embryonic day (E) 13.5 before collection at E17.5. Successful electroporation was confirmed by immunohistochemistry using established methods (41). Primary antibodies were chicken anti-GFP 1/1000 (Abcam ab13970) and rabbit anti-RFP 1/500 (Abcam ab62341), and secondary antibodies coupled to Alexa-488 or Alexa-546 1/200 (Invitrogen). Nuclei were stained with DAPI. Images were acquired on a Leica SP8 confocal microscope and processed using ImageJ. Sample brains, 4 CHD8 TaDa and 3 Dam-only, were dissected and frozen for library processing. All mouse husbandry and experiments were carried out in a Home Office-designated facility, according to the UK Home Office guidelines upon approval by the local ethics committee (project license PPL70/8727).

Targeted DamID-seq (TaDa-seq) libraries were prepared as previously described (27). Sample genomic DNA extraction was performed using the Qiagen QIAamp DNA Micro Kit (Qiagen, 56304). Extracted genomic DNA was digested overnight at 37°C with DpnI (NEB, R0176S) to cut adenine-methylated GATC sites. Following digestion, DNA was column purified with the QIAquick PCR Purification Kit (Qiagen, 28104) to remove un-cut genomic DNA. dsADR adaptors were blunt-end ligated to DpnI-digested fragments using T4 DNA ligase (NEB, M0202S; 2 hours at 16°C, heat inactivation at 65°C for 20 minutes) to prepare for PCR amplification. Before PCR amplification, fragments were digested with DpnII (NEB, R0543S) to cut non-methylated GATC sites and prevent amplification of unmethylated regions and purified with a 1:1.5 ratio of Seramag beads (Fisher Scientific, 65152105050250). PCR amplification of DpnII-digested fragments using MyTaq (Bioline, BIO-21112) enriched for methylated fragments before samples were sonicated and prepped for sequencing. Sonicated samples were subjected to AlwI digestion (NEB, R0513S) to remove previously ligated adaptors and initial GATC sequences from fragments. A modified TruSeq protocol was used to generate sequencing libraries involving end repair, 3’ end adenylation, sequencing adaptor ligation, and DNA fragment enrichment using a reduced number of PCR cycles. TaDa-seq libraries were sequenced on the Illumina HiSeq 1500 platform using a single-end 50bp strategy by the Gurdon Institute Next Generation Sequencing Core.

### Computational analysis of TaDa-seq and ChIP-seq Datasets

Sequenced TaDa-seq libraries were analyzed to identify genomic regions with enriched coverage for CHD8 TaDa and Dam-only libraries. Representative CHD8 ChIP-seq datasets were downloaded from the Sequence Read Archive (19, GSE57369; 22, PRJNA379430). Unaligned TaDa-seq and ChIP-seq reads were trimmed using TrimGalore (Version 0.4.2), assessed for general quality control with the FastQC tool (Version 0.11.9), and aligned to the mouse reference genome (mm10) using BWA (Version 0.7.17). Biological replicates were analyzed independently and as a single merged file generated via samtools (Version 1.10). Coverage plots were generated independently for Dam-only and CHD8 TaDa-seq replicates (deepTools, RPKM normalization). Merged CHD8 TaDa-seq was normalized against Dam-only (33) for visualization of coverage and enrichment. TaDa-seq peak calling was performed using MACS2 (Version 2.2.5) with model-based peak identification disabled, a p-value cutoff set at less than 0.00001, and the merged Dam-only dataset as a control. Peak calling for individual CHD8 TaDa-seq replicates was performed against the merged Dam-only dataset to identify specific peaks that were enriched in CHD8 TaDa versus non-specific signal in Dam-only experiments. Peak calling for the Dam-only merged dataset was performed without a control dataset as Dam-only is analogous to assays of accessible chromatin. Peak calling for CHD8 ChIP-seq experiments was performed using the same MACS2 parameters, including comparison to input controls. A final set of merged CHD8 TaDa-seq peaks was obtained using bedtools intersect (Version 2.29.2) to select high confidence peaks that were present in at least 3 replicates and had a MACS2 FDR less than 0.00001.

Enriched regions from TaDa-seq and ChIP-seq datasets were annotated to genomic features using custom R scripts and combined UCSC and RefSeq transcript sets (25). CHD8 target genes were assigned to nearest transcription start site, which for distal peaks was achieved using the bedtools closest command (Version 2.29.2). Bigwig coverage files were generated using deeptools bamCoverage (Version 3.3.1). Embryonic E16.5 bigwig coverage files from the ENCODE Consortium portal (42, https://www.encodeproject.org/) were downloaded to compare CHD8 datasets with open chromatin and histone marks (Experiments: ENCSR428OEK, ENCSR658BBG, ENCSR587JRQ, ENCSR141ZQF, ENCSR836PUC, ENCSR129DIK). Genome-wide signal summary Spearman correlation heatmaps using the default bin size of 10 kb were generated using the multiBigwigSummary and plotCorrelation tools from deeptools (Version 3.3.1). Differences in signal intensity between CHD8 TaDa-seq replicates in the correlation heatmaps were due to differences in sequencing depth. Peak loci heatmaps were generated using the deeptools computeMatrix and plotHeatmap tools (Version 3.3.1). Intersection of called peaks was performed using bedtools intersect (Version 2.29.2) with CHD8 TaDa-seq filtered peaks and ChIP-seq datasets. Promoter-proximal versus promoter-distal and peak set concordance datasets were also obtained using the bedtools intersect tool (Version 2.29.2). Ontology analysis was performed using the GREAT online tool (43, Version 4.0.4) or goseq (44, Version 1.36.0). HOMER was used to perform de novo motif discovery with default parameters (45, Version 4.10). Comparison between CHD8 TaDa-seq and E17.5 RNA-seq data was performed using previously published RNA-seq data (21, 25). Data that support the findings of this study are available from GEO (Accession #TBD). Genomic coverage datasets are available as Track Hubs for visualization using the UCSC Genome Browser and all analysis scripts are available at https://github.com/NordNeurogenomicsLab/.

## RESULTS

### Cloning and *in utero* electroporation of CHD8 TaDa plasmid into embryonic mouse cortex

To study CHD8 binding patterns in embryonic neurodevelopment, we used an established Targeted DamID-seq (TaDa-seq) protocol (27), combined with *in utero* electroporation of embryonic day (E) 13.5 mouse brain (Figure 1A-D). A full-length human CHD8 ORF was cloned into the TaDa construct and sequence verified. The experimental TaDa constructs (CHD8 TaDa and Dam-only) are designed to express extremely low levels of the CHD8-Dam fusion proteins (Figure 1A; Methods). Recruitment of CHD8 TaDa to specific genomic loci, either directly or through interaction with other proteins, will be detected by TaDa (Figure 1E), while Dam-only will interact non-specifically genome-wide at regions of accessible chromatin. The CHD8 TaDa and Dam-only signals are compared to distinguish CHD8-specific interactions from generally accessible chromatin (Figure 1E).

**Figure 1.**
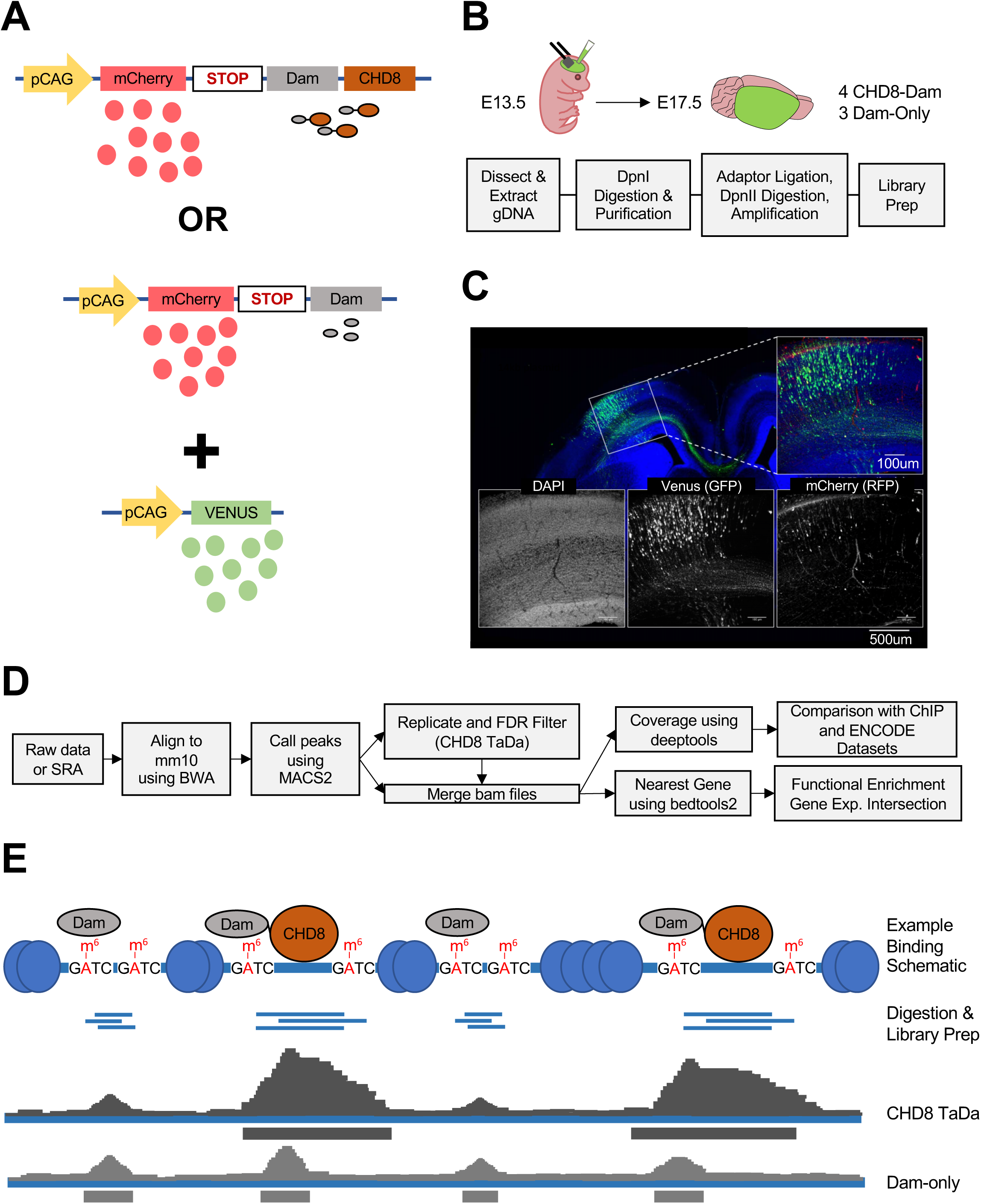
Overview of the Targeted DamID followed by sequencing (TaDa-seq) pipeline. (**A**) Plasmids used for TaDa. The top plasmid is a diagram for CHD8 TaDa experiments. The middle plasmid is a diagram for Dam-only experiments. The bottom plasmid is a diagram for the *in utero* electroporation control injected with the CHD8 TaDa or Dam-only plasmids. (**B**) Schematic and flowchart of TaDa-seq experiments. E13.5 mouse embryos were injected with CHD8 TaDa or Dam-only plasmid and the *in utero* electroporation control plasmid. Four CHD8 TaDa and three Dam-only brains from the same litter were dissected. Frozen brains were then processed for the pipeline indicated in the grey boxes. (**C**) Immunohistochemistry showing overlap between green fluorescence (in utero electroporation control), red fluorescence (mCherry expression upstream of the CHD8 TaDa open reading frame), and DAPI (nuclei) illustrates successful transfection of experimental plasmids. (**D**) TaDa-seq computational analysis pipeline used in this study. (**E**) Schematic showing example signal from CHD8 TaDa or Dam-only protein binding at genomic loci.

The Venus control and CHD8 TaDa or Dam-only constructs were electroporated *in utero* into developing mouse cerebral cortex at E13.5 (Figure 1B). A pCAG-Venus construct was co-electroporated with CHD8 TaDa and Dam-only plasmids as a delivery control to visualize the electroporated region. Following delivery, there was a 4-day period where the constructs could be expressed in cells that took up the plasmids before tissues were collected at E17.5. Representative images of E17.5 cortex show green immunofluorescence representing Venus expression confirming *in utero* electroporation into developing somatosensory cortex, while red immunofluorescence shows expression of the primary open reading frame of the TaDa construct, mCherry (Figure 1C). Translation of the CHD8 TaDa or Dam-only open reading frames is too low to detect by immunostaining. Following IUE, transfected radial glial neural progenitor cells undergo self-renewal as well as producing early neurons that will migrate to form the layers of the cortex. The incubation period represented the window during which Dam methylation occurred, resulting in Dam activity from ventricular zone progenitors to early cortical neurons, evidenced by Venus and mCherry expression (Figure 1C).

### Genomic patterns of CHD8 TaDa-seq and representative CHD8 ChIP-seq datasets

For sequence-based analysis, we collected 4 CHD8 TaDa and 3 Dam-only samples from the same litter of MF-1 outbred mice and processed them using the TaDa-seq experimental and computational pipeline (27, 33; see Methods for details, Figure 1B-D). Individual replicates and merged datasets for CHD8 TaDa-seq and Dam-only experiments were analyzed. Coverage plots were generated to show signal independently in the CHD8 TaDa-seq and Dam-only datasets, and CHD8 TaDa-seq coverage was additionally normalized using Dam-only to visualize enrichment representing CHD8-specific interactions. Enriched genomic regions identified in at least three of four CHD8 TaDa-seq replicates at high statistical stringency were considered to be high confidence CHD8 interaction regions. In addition to serving as a non-specific control for CHD8 TaDa, regions identified as enriched in Dam-only experiments are expected to represent accessible chromatin (46). Following peak calling and merging, there were 142,375 enriched peaks in the Dam-only experiments and 24,533 that passed stringent significance and reproducibility criteria across CHD8 TaDa-seq experiments.

To verify specificity and relevance of CHD8 targets mapped by TaDa-seq, we compared the CHD8 TaDa-seq dataset to two published CHD8 ChIP-seq experiments performed on mouse brain. The first dataset was time- and tissue-matched with our TaDa-seq data, profiling E17.5 mouse cortex (19). The second was from adult mouse cortex (22). While CHD8 ChIP-seq datasets vary in results, these ChIP-seq datasets had no evidence of technical issues and were highly correlated to each other and other CHD8 ChIP-seq datasets (25). Raw sequence files were downloaded and analyzed using standard approaches to generate coverage and peak intervals (25; see Methods), with 44,383 and 32,335 peaks mapped in the E17.5 and adult cortex datasets, respectively.

We examined patterns of enrichment between CHD8 TaDa-seq, CHD8 ChIP-seq, and epigenomic datasets generated for E16.5 mouse cortex via ENCODE. First, we examined genomic loci that were previously found to have consistent and strong CHD8 peaks across ChIP-seq datasets (Figure 2, Supplementary Figure 2). For example, promoter interactions for genes associated with RNA processing, such as *Hnrnpll, Srsf7, Srsf1*, and *Sf3b1*, or genes associated with chromatin remodeling, such as *Top1* (Figure 2A-B). Read coverage at these loci illustrates reproducibility and specificity of CHD8 TaDa protein genomic interactions as compared with chromatin accessibility revealed by Dam-only (Figure 2A). As expected, Dam-only peaks occurred throughout the genome, indicating expected non-specific adenine methylation in regions of accessible chromatin. Comparison of CHD8 genome interactions identified by TaDa-seq with the two published CHD8 ChIP-seq datasets showed strong concordance in enrichment at these loci, indicating TaDa-seq captured reproducible interactions between CHD8 and its genomic targets.

**Figure 2.**
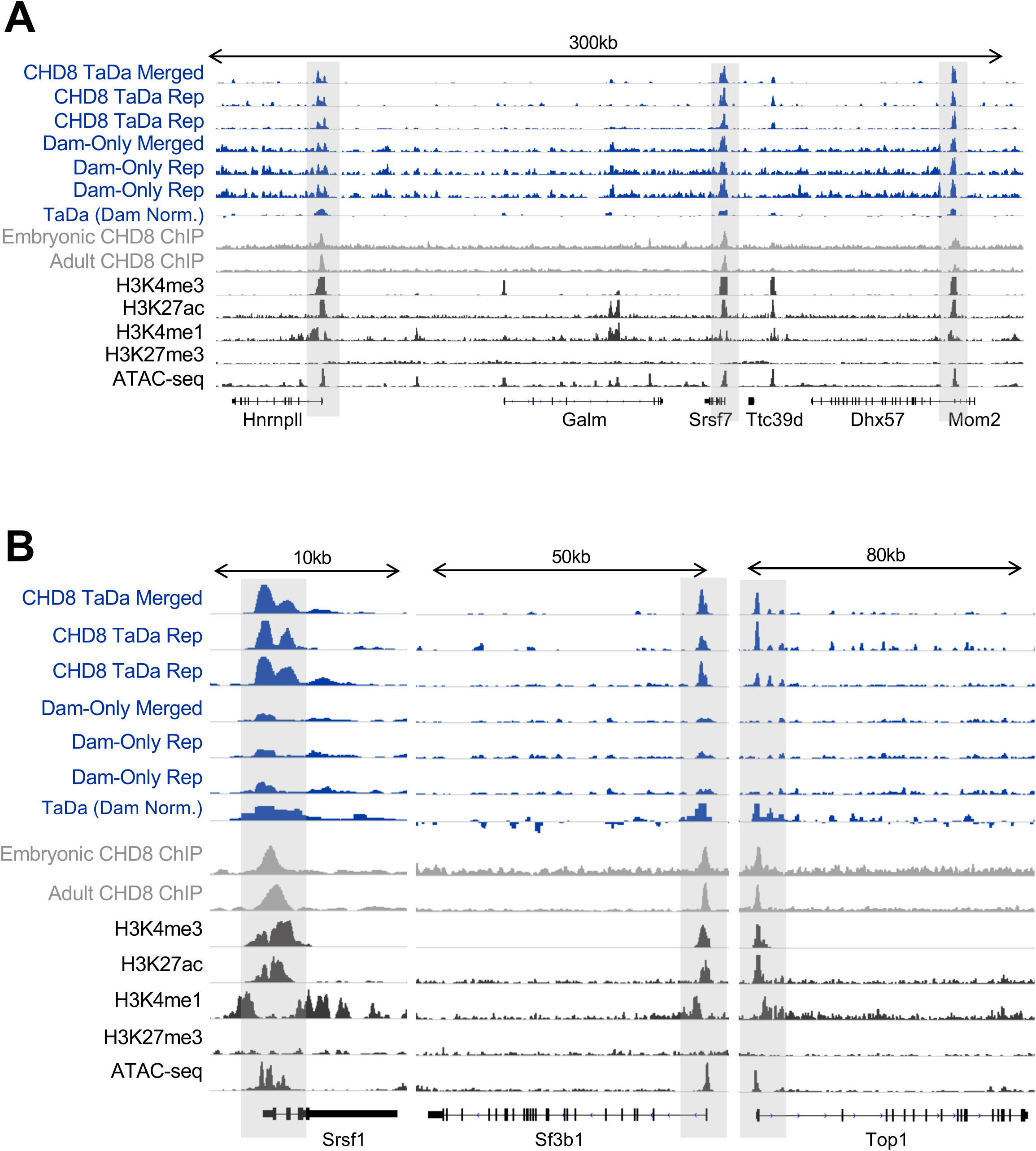
Recapitulation of CHD8 binding near promoters across the genome. Data showing CHD8 binding at loci previously identified in CHD8 binding characterization studies, including RNA processing genes, *Hnrnpll* and *Srsf7* in panel A, *Srsf1* and *Sf3b1* in panel B, and a chromatin remodeling gene, *Top1*, in panel B. Grey boxes highlight CHD8 binding near identified promoters of interest. CHD8 TaDa, Dam-only, or Dam-only normalized CHD8 TaDa (TaDa Dam Norm.) experiment tracks are in blue (representative biological replicates shown), CHD8 ChIP-seq experiments are in grey, and datasets of histone and chromatin accessibility signatures from the ENCODE consortium are in black. Linear representations of genes from the mouse mm10 genome are shown below coverage tracks. Height of the y-axis is scaled to show the peak for each track separately.

We compared our datasets for CHD8 TaDa, Dam-only, and CHD8 ChIP-seq (Figure 3A-C) and plotted coverage heatmaps for comparison of signal enrichment between CHD8 TaDa, Dam-only, and CHD8 ChIP-seq datasets (Figure 3B). CHD8 TaDa-seq and CHD8 ChIP-seq both found enrichment of CHD8 binding strongly enriched at promoters, though distal interactions were also present (25, Figure 3A). Consistent with the individual loci in Figure 2, CHD8 TaDa-seq enrichment was strongly correlated with CHD8 ChIP-seq signal. This observation, coupled with reduced CHD8 TaDa-seq enrichment for the majority of Dam-only peaks, confirmed the specificity of CHD8 TaDa binding throughout the genome (Figure 3B). No single DNA motif was identified at CHD8 target loci defined by either TaDa-seq or ChIP-seq, suggesting CHD8 interactions were not generally guided by direct binding to a recognition sequence (Supplementary Figure S1). These data show results from CHD8 TaDa-seq experiments are consistent with previous ChIP-seq observations of CHD8 genomic binding activity. While many of the same loci were captured, enrichment rank varied between CHD8 TaDa-seq and ChIP-seq. Most peaks that were called in only the CHD8 TaDa-seq or the ChIP-seq datasets exhibited sub-significant signal in the other assay (Figure 3B-C). This suggests differences in interaction targets between CHD8 TaDa-seq and ChIP-seq was largely due to sensitivity of detection and peak calling stringency.

**Figure 3.**
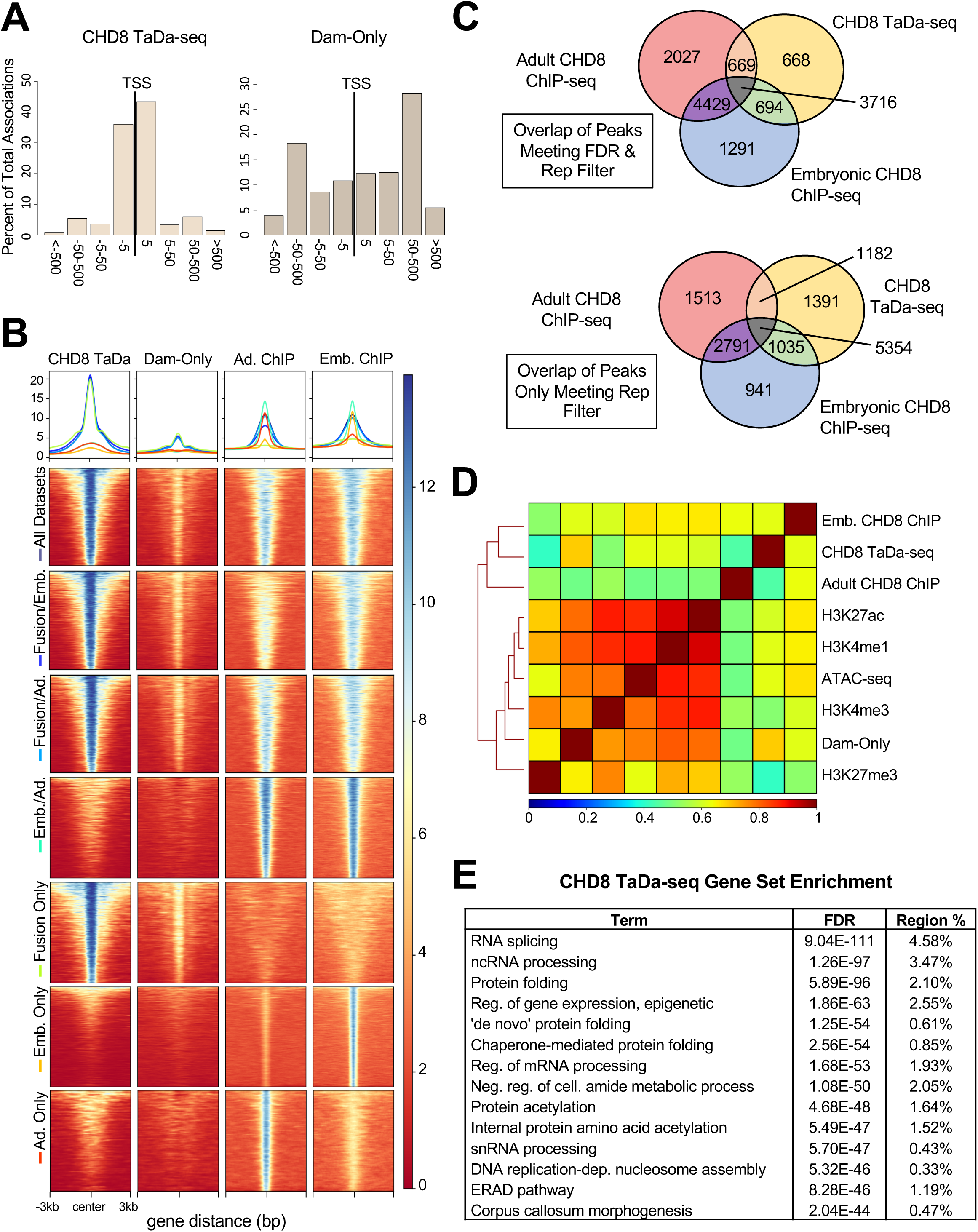
Computational Comparison of CHD8 binding shows correspondence in signal across TaDa-seq and ChIP-seq experiments. (**A**) Bar plots showing association of peaks with transcription start sites (TSS) using the GREAT online analysis tool. Bins along the x-axis represent 5, 50, 500, and greater than 500 kilobases away from the nearest TSS. (**B**) Genome-wide coverage heatmaps showing enrichment of signal at peaks for each dataset indicated on the left-hand side. Y axis of datasets were matched for visual comparison. Small line plots indicate the average normalized peak enrichment for each dataset with the color for each line next to each dataset name. Each peak is centered along the middle of each plot with a 3 kilobase pair window on each side. The legend indicates normalized enrichment. (**C**) Venn diagram showing the number of peaks annotated to genes overlapping with CHD8 TaDa, Embryonic CHD8 ChIP-seq, and Adult CHD8 ChIP-seq using stringent CHD8 TaDa-seq peak thresholding with peaks meeting an FDR < 0.00001 cutoff in at least 3 replicates (Top) or a looser threshold of peaks present in at least 3 replicates (Bottom). (**D**) Genome coverage correlation heatmap showing relationship between representative CHD8 TaDa-seq, Dam-only, CHD8 ChIP-seq, and ENCODE histone mark and chromatin accessibility datasets. Data are hierarchically clustered according to genome-wide similarity as indicated by a dendrogram. Legend indicates the correlation value between datasets. H3K27me3 is a histone mark associated with repressed DNA loci. H3K4me3 is a histone mark associated with actively transcribed promoters. ATAC-seq is sequencing data of open chromatin regions. H3K4me1 and H3K27ac are histone marks associated with putative enhancers. (**E**) Table showing functional annotations associated with CHD8 TaDa-seq called peaks. Region % refers to the percent of the total peak set annotated to each term.

As predicted, the Dam-only genome-wide signal strongly correlated with ENCODE E16.5 fetal cortex ATAC-seq datasets, confirming that Dam-only binding is enriched at accessible chromatin. Dam-only datasets were also strongly correlated with H3K4me3, H3K4me1, and H3K27ac marks, consistent with the relationship between accessible chromatin and transcriptionally active chromatin states at promoters and enhancers. Genome-wide, quantitative signal between CHD8 TaDa-seq and ChIP-seq datasets were moderately correlated (Figure 3D, Supplementary Figure S3), consistent with differences in loci enrichment strength but similar interaction targets between the methods as previously shown with other targets (37). The CHD8 TaDa-seq and matched E17.5 cortex CHD8 ChIP-seq datasets were also strongly correlated with ATAC-seq and histone marks associated with open and transcriptionally active chromatin, H3K4me3/me1 and H3K27ac. CHD8 TaDa-seq and ChIP-seq datasets showed reduced correlation with a mark for repressive chromatin, H3K27me3. Overall, these results support for a primary role of CHD8 in transcriptional activation in embryonic mouse cortex.

### CHD8 TaDa-seq indicates direct role of CHD8 in transcriptional activation of genes associated with cellular homeostasis and novel distal enrichment at neuronal loci

Gene ontology analysis using GREAT (43) showed CHD8 TaDa-seq peaks had the strongest enrichment for genes associated with general cellular homeostasis, and specifically with RNA splicing, protein folding, and chromatin regulation genes (Figure 3E, Supplementary Table 1). This finding is consistent with previous findings of CHD8 ChIP-seq datasets (25). There was also evidence for reduced, but still significant enrichment of CHD8 interactions at loci associated with metabolism and neuron differentiation, also in line with earlier evidence. Intersection of genes associated with CHD8 TaDa-seq peaks here with transcriptomic data from a published analysis of E17.5 cortex from mice harboring heterozygous *Chd8* mutations (21) showed that genes associated with strong CHD8 TaDa-seq peaks are both highly expressed in E17.5 embryonic mouse cortex and more likely to be downregulated as a consequence of *Chd8* haploinsufficiency (Figure 4A, 4C). CHD8 TaDa-seq interactions were less likely to be found at loci that were upregulated as a consequence of *Chd8* haploinsufficiency (Figure 4D). There was no enrichment for Dam-only interactions for downregulated genes (Figure 4B). Thus, CHD8 TaDa-seq results provide evidence for CHD8-dependent activation of highly expressed genes associated with general cellular functions, consistent with results from individual CHD8 studies and from meta-analysis of published CHD8 ChIP-seq data (18–25).

**Figure 4.**
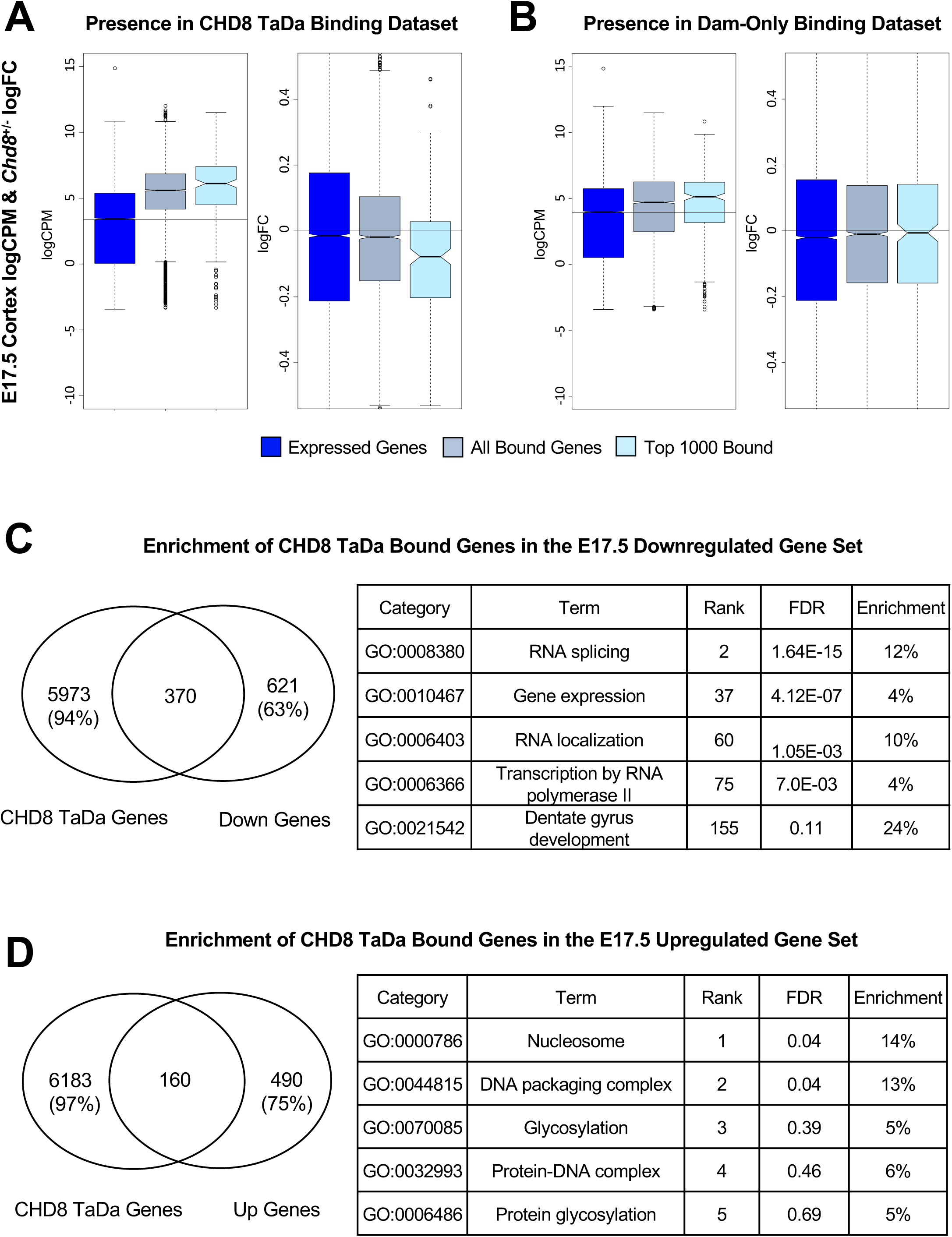
CHD8 Binding is Associated with Activation of Highly Expressed Genes. (**A**) Box and whisker plots showing comparison between CHD8 TaDa-seq peak rank and an E17.5 *Chd8* haploinsufficiency differential gene expression dataset. Change in log fold counts per million of genes according to CHD8 binding (Left). Change in log fold change of genes according to CHD8 binding (Right). Boxes were plotted according to CHD8 binding affinity bins: all genes meeting at least 0.1 count per million sequencing coverage (Expressed Genes), any genes having CHD8 binding (All Bound Genes), and the top 1000 genes near CHD8 peaks (Top 1000 Bound). Notches indicate values within the 95% confidence interval of the median. (**B**) Box and whisker plots showing comparison between Dam-only peak rank and an E17.5 *Chd8* haploinsufficiency differential gene expression dataset. Change in log fold counts per million of genes according to Dam binding (Left). Change in log fold change of genes according to Dam binding (Right). Boxes were plotted according to CHD8 binding affinity bins: all genes meeting at least 0.1 count per million sequencing coverage (Expressed Genes), any genes having CHD8 binding (All Bound Genes), and top 1000 genes near CHD8 peaks (Top 1000 Bound). Notches indicate values within the 95% confidence interval of the median. (**C-D left**) Venn diagrams indicating the number of genes overlapping between the CHD8 TaDa-seq and E17.5 *Chd8* haploinsufficiency significant (p < 0.05) downregulated and upregulated datasets. (**C-D right**) Tables showing functional annotations associated with genes having CHD8 binding in downregulated (C) and upregulated (D) genes from the E17.5 *Chd8* haploinsufficiency dataset (p < 0.05) using goseq. Enrichment values indicate the percent of genes in the dataset that are differentially expressed and bound by CHD8 via TaDa-seq in relation to the total number of genes associated with each term.

While CHD8 TaDa-seq defined interactions were strongly enriched at promoters, some loci also exhibited distal peaks. Loci with CHD8 TaDa-seq distal enrichment, or enrichment outside of promoters, were more strongly enriched for genes associated with neurodevelopmental and neuron-specific function. This suggests distinct proximal versus distal targets for CHD8 in embryonic mouse cortex. Examples of neuronal genes with distal CHD8 interactions include genes with dual roles in gene regulation and neurodevelopment, such as *Myt1l* (Figure 5A), as well as genes having more specific roles in neuronal morphology and synaptic signaling, such as *Ank3* and *Dlg4*, which encodes PSD95, (Figure 5B-C). Comparison of these loci with marks for putative enhancers (H3K27ac, H3K4me1), open chromatin (ATAC), and transcriptional activation (H3K4me3) suggests that CHD8 TaDa-seq distal peaks intersect with distal cis-regulatory sequences. The distal CHD8 interactions identified in our CHD TaDa-seq data are somewhat captured by the representative E17.5 CHD8 ChIP-seq datasets, but with reduced enrichment relative to the TaDa-seq signal (Figure 5A-C). To assess whether gene sets bound by CHD8 at their promoters and those targeted distally indeed are enriched for different functional categories, we split CHD8 TaDa-seq peaks into promoter proximal (within 1 kb of TSS) and promoter distal interactions. Loci with promoter binding mirrored the overall analysis (Figure 5D). Distal CHD8 interactions were also enriched at loci associated with general regulatory function terms such as “negative regulation of transcription” and “negative regulation of RNA metabolic process.” However, loci associated with distal CHD8 interactions identified via TaDa-seq were more strongly enriched for brain development and neuronal functions, for example “cell morphogenesis involved in neuron differentiation,” “regulation of dendritic spine development,” and “cell fate commitment” (Figure 5D, Supplementary Table 1). CHD8 ChIP-seq datasets similarly split into proximal and distal interactions suggest concordant patterns, though with substantially reduced enrichment (Supplementary Figure S4, Supplementary Table 1). This difference in CHD8 distal interaction profile appeared to be the most relevant distinction between results from CHD8 TaDa-seq and the representative CHD8 ChIP-seq datasets.

**Figure 5.**
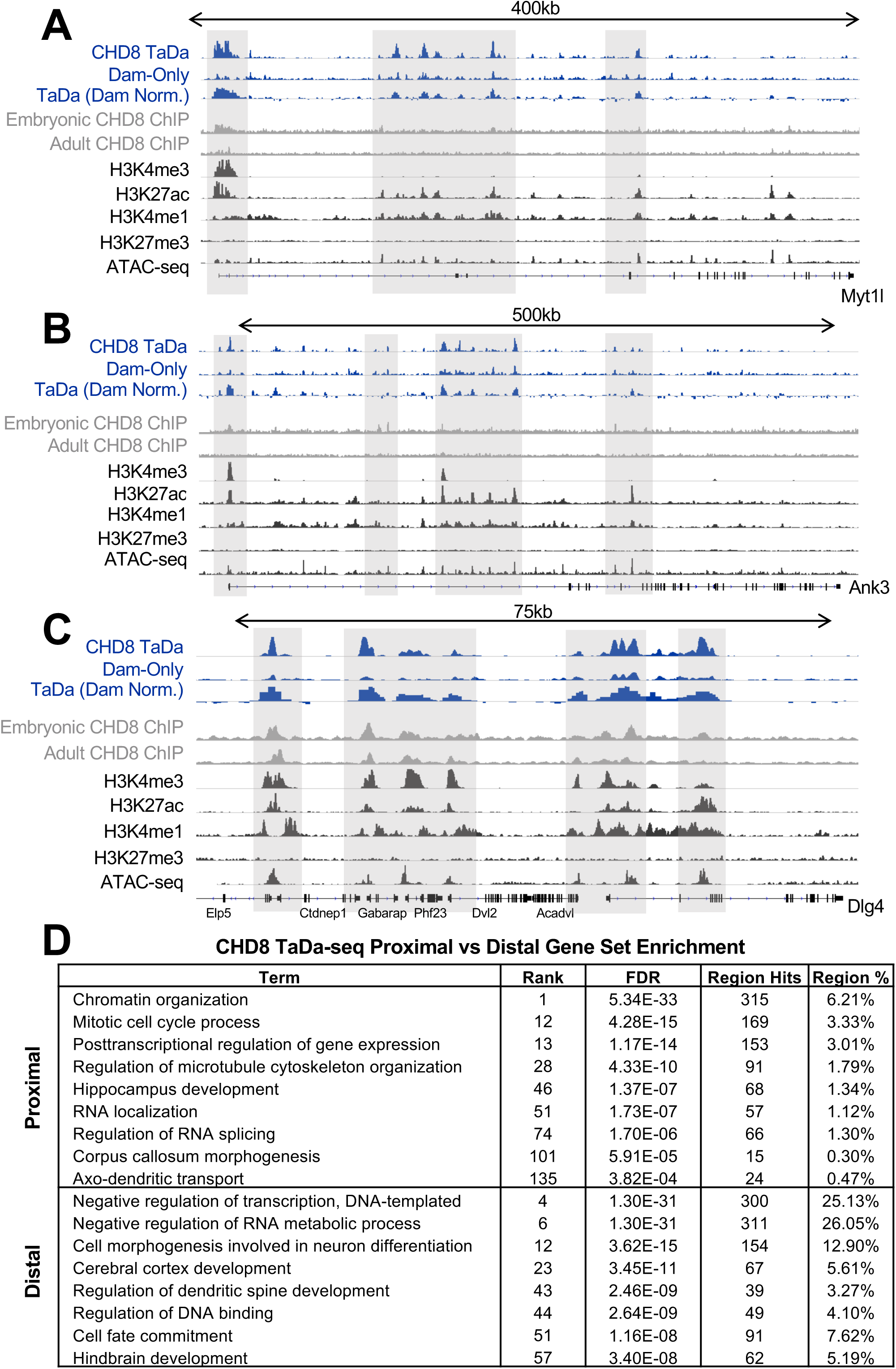
TaDa-seq Identifies Both Promoter Proximal and Promoter Distal CHD8 binding. (**A-C**) CHD8 binding near genes important for regulation of neuronal gene expression, *Myt1l* (A), and synaptic function, *Ank3* (B) and *Dlg4* (C). Grey boxes highlight CHD8 binding near select promoter and distal regions of interest overlapping with putative enhancer marks (H3K27ac and H3K4me1). CHD8 TaDa-seq experiment tracks are in blue, CHD8 ChIP-seq experiments are in grey, and datasets of histone and chromatin accessibility signatures from the ENCODE consortium are in black. Linear representations of genes from the mouse mm10 genome are shown below coverage tracks. Height of the y-axis is scaled to show the peak for each track separately. (**D**) Table showing functional annotations associated with promoter proximal (<1kb from TSS) (Top) and promoter distal (Bottom) regions. Rank refers to the rank within the dataset. A rank of 1 would mean the annotation with the smallest FDR value (aka the most significant). Region hits capture the number of peaks associated with each term. Region % captures the percent of regions captured compared to the total number of peaks.

## DISCUSSION

We successfully implemented Targeted DamID followed by sequencing (TaDa-seq) *in vivo* in embryonic mouse brain by *in utero* electroporation, characterizing genomic interactions of the NDD-relevant chromatin remodeler, CHD8. We chose to study a chromatin remodeling protein where ChIP-seq grade antibodies were available and ChIP-seq had been repeatedly performed, enabling comparison of TaDa-seq and ChIP-seq. While CHD8 ChIP-seq studies have provided valuable insights regarding CHD8 molecular function *in vivo* in mouse brain and across *in vitro* models, variable results across individual ChIP-seq experiments can confound interpretations of CHD8 activity and gene regulation. Thus, our study represents a proof-of-principle implementation of TaDa-seq in the context of *in vivo* mouse brain development and advances understanding of *CHD8*, a leading NDD risk gene. These findings open new avenues to interrogate the function of proteins that are intractable or technically challenging to study using ChIP-seq. The interactions identified here using TaDa-seq have orthogonally validated CHD8 interactions mapped using ChIP-seq and highlight the presence of CHD8 distal interactions at NDD-relevant neurodevelopmental and neuronal genes.

Implementation of TaDa-seq requires up-front steps of construct generation and delivery for expression in the cells or tissues of interest. Furthermore, the Dam methylase must be expressed and have time to methylate at genomic sites. In contrast, ChIP-seq can be performed on unmodified cells or tissues and captures interactions present at a specific time. However, ChIP-seq grade antibodies must be available, crosslinking and fragmentation is generally required, and protein-DNA interactions must be strong enough to enable sensitive capture using immunoprecipitation. Furthermore, TaDa-seq requires substantially less material than typical ChIP-seq methods (35, 37). Thus, while ChIP-seq remains a more generally applicable method with clear temporal resolution, we show that TaDa-seq can overcome barriers that negatively-impact ChIP-seq performance and that TaDa-seq can capture protein-DNA interactions that might be missed due to sensitivity thresholds of ChIP-seq. Conditional expression of the TaDa-seq constructs with cell-type specific promoters would enable identification of cell-type-specific chromatin interactions, as has been shown in *Drosophila* (26, 34, 36, 38). Such an approach offers the potential to address key questions regarding context specific function and genomic interactions of chromatin remodelers and other DNA-associated proteins in the developing mouse brain.

By directly comparing CHD8 TaDa-seq and ChIP-seq, we found that TaDa-seq experiments are highly reproducible and perform well with regard to sensitivity and specificity, with strong overall concordance between the interactions mapped by these different methods. TaDa-seq thus joins the few published methods for resolving protein-DNA interactions genome-wide that can be deployed *in vivo* and do not require cross-linking or immunoprecipitation. Recently, another such method mapped transcription factor interactions at single cell resolution by fusing transcription factors to transposase domains and locating transposition events through direct DNA sequencing (47). Application of TaDa-seq offers the opportunity to characterize the neurodevelopmental function of chromatin remodeler proteins implicated in NDDs, as we did here for *CHD8*, that might be difficult to interrogate using ChIP-seq. For example, TaDa-seq has been used to map the binding sites of kismet, the Drosophila ortholog for CHD8, which has roles in cell proliferation, synaptic transmission, axonal pruning, circadian rhythm, and memory (48).

Overlapping sets of CHD8 interactions were largely captured by both TaDa-seq and ChIP-seq technologies, but with differences in signal strength. This could be due to general differences in performance, or CHD8-specific features due to strong correlation between CHD8 binding and open chromatin at promoters. Previous comparisons of TaDa-seq and ChIP-seq *in vitro* and in *Drosophila* have found similar evidence for general concordance in target loci but divergence in quantitative strength (26–27, 33–35, 37). In our study, loci identified in ChIP-seq but not TaDa-seq, appeared to represent a random sampling of loci with weaker ChIP-seq signal and sub-significant CHD8 TaDa-seq signal. Similarly, loci that were significant in CHD8 TaDa-seq but not ChIP-seq exhibited weak enrichment in ChIP-seq datasets. It is possible that ChIP-seq may have reduced sensitivity for the distal interactions that were preferentially captured by TaDa-seq due to reduced interaction stability or the transient nature of these interactions compared to very strong CHD8 promoter interactions. By using adenine methylation as the readout, TaDa-seq does not seem to be as impacted by over-sampling of stronger protein-DNA interactions, and may thus better capture transient or weaker interactions. Alternatively, as TaDa-seq captures adenine methylation throughout the incubation time of E13.5 to E17.5, it is possible that some of the TaDa-seq specific CHD8 interactions are limited to stages earlier than profiled via ChIP-seq at E17.5. In summary, CHD8 interactions detected by TaDa-seq and ChIP-seq were overall highly concordant, with some evidence for assay-specific differences such that the combined interaction sets can complement each other.

Consistent with previous conclusions, our study confirms that the majority of CHD8 interactions occur at promoters. There was no evidence for direct binding of CHD8 to a primary DNA motif, supporting a model of CHD8 recruitment by co-factors or transcription factors. Our results also support a direct role in transcriptional activation. CHD8 interactions were strongly correlated with open chromatin assayed by ATAC-seq and with histone marks associated with open and actively transcribed promoters. Loci with CHD8 interactions were also more likely to be downregulated due to *Chd8* haploinsufficiency. Finally, comparison between CHD8 TaDa, Dam-only, and ENCODE data clearly showed that CHD8 interactions are specific to a subset of promoter and distal loci, rather than broadly co-occurring with accessible chromatin or with the global deposition of any specific histone modifications. Our CHD8 TaDa-seq results further establish the strong enrichment of CHD8 binding near promoters of genes associated with general cellular functions involved in replication, chromatin, transcription, and translation. The TaDa-seq data also highlight potential CHD8 involvement in distal regulation for a subset of neurodevelopmental and neuronal genes.

The unexpected increased significance for CHD8 binding near distal regions in the TaDa-seq experiments indicates that using orthogonal approaches to ChIP-seq may bring novel insights due to differing detection biases. The evidence here for a brain-specific CHD8 distal interaction signature has significant potential implications for models of the role of CHD8 in brain development and function. These distal regions overlap with H3K27ac, a histone mark associated with putative enhancers, suggesting a role related to distal regulatory elements. It is possible that CHD8 is involved in distal chromatin remodeling or enhancer activation in the developing brain, parallel to what has been reported in the context of CHD8 in estrogen response (18). While *CHD8* is an essential gene, there are opposite effects on cortex development between *Chd8* null knockout mice, exhibiting microcephaly, and mice heterozygous for a *Chd8* mutation, exhibiting macrocephaly. It is possible that *CHD8* haploinsufficiency has a specific effect on a subset of CHD8 interactions, for example disrupting weaker interactions. Further studies are needed to explore the difference in CHD8 function in the brain at promoters versus distal sites and dosage sensitivity of these interactions in the context of *CHD8* haploinsufficiency. Future studies are also necessary to determine the context-specific protein interaction partners of CHD8 to understand its role in transcriptional regulation in the brain.

In summary, this study shows the value of TaDa-seq as an alternative to ChIP-seq, with the novel implementation to map protein-DNA interactions in embryonic mouse cortex. Implementation of CHD8 TaDa-seq revealed a comprehensive set of CHD8 target loci in the genome, furthering understanding of the genomic function of CHD8 in the developing brain and the relationship between CHD8 interaction targets and ASD-and ID-relevant pathology caused by *CHD8* mutations. This work serves as a model for studying other proteins, including the many chromatin remodeling factors associated with NDDs for which ChIP-seq may be technically challenging or where ChIP-seq grade antibodies unavailable.

## DATA AVAILABILITY

Data that support the findings of this study are available from GEO (Accession #TBD) or upon request.

## Author contributions

A.A.W, A.S.N., and A.H.B. conceived of the project. J.v.d.A., S.W.C. and R.Y. performed experiments. A.A.W. performed computational analysis. A.A.W. and A.S.N. drafted the manuscript. All authors contributed to manuscript revisions.

## FIGURES

**Supplementary Figure S1.**
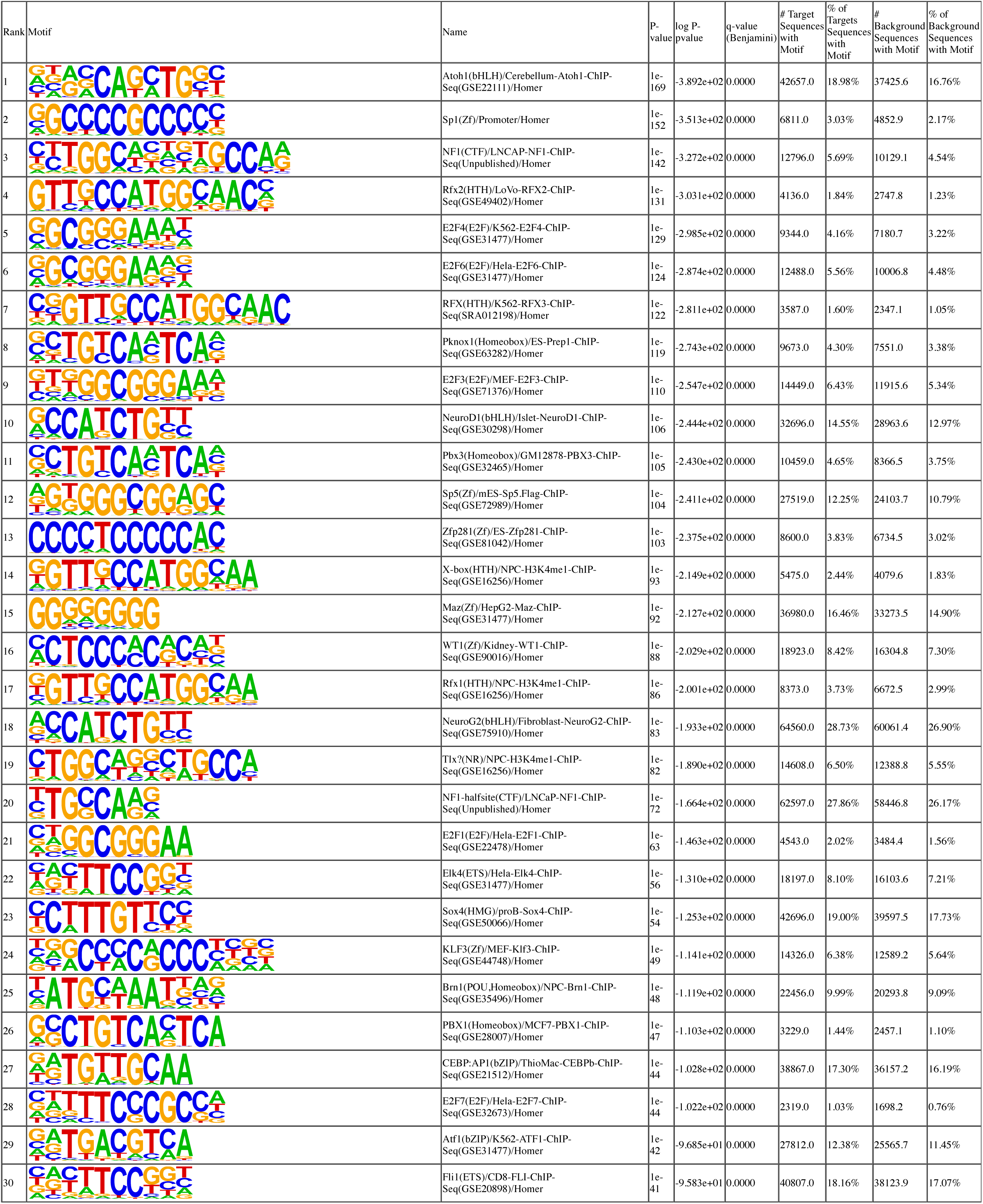
CHD8 Binding is not directed by a single specific motif. HOMER results of the top 30 motifs identified in CHD8 TaDa-seq peaks. Target sequences refer to sequences from CHD8 TaDa-seq peaks used as input. Background sequences refers to random sequence-content matched intervals in the genome.

**Supplementary Figure S2.**
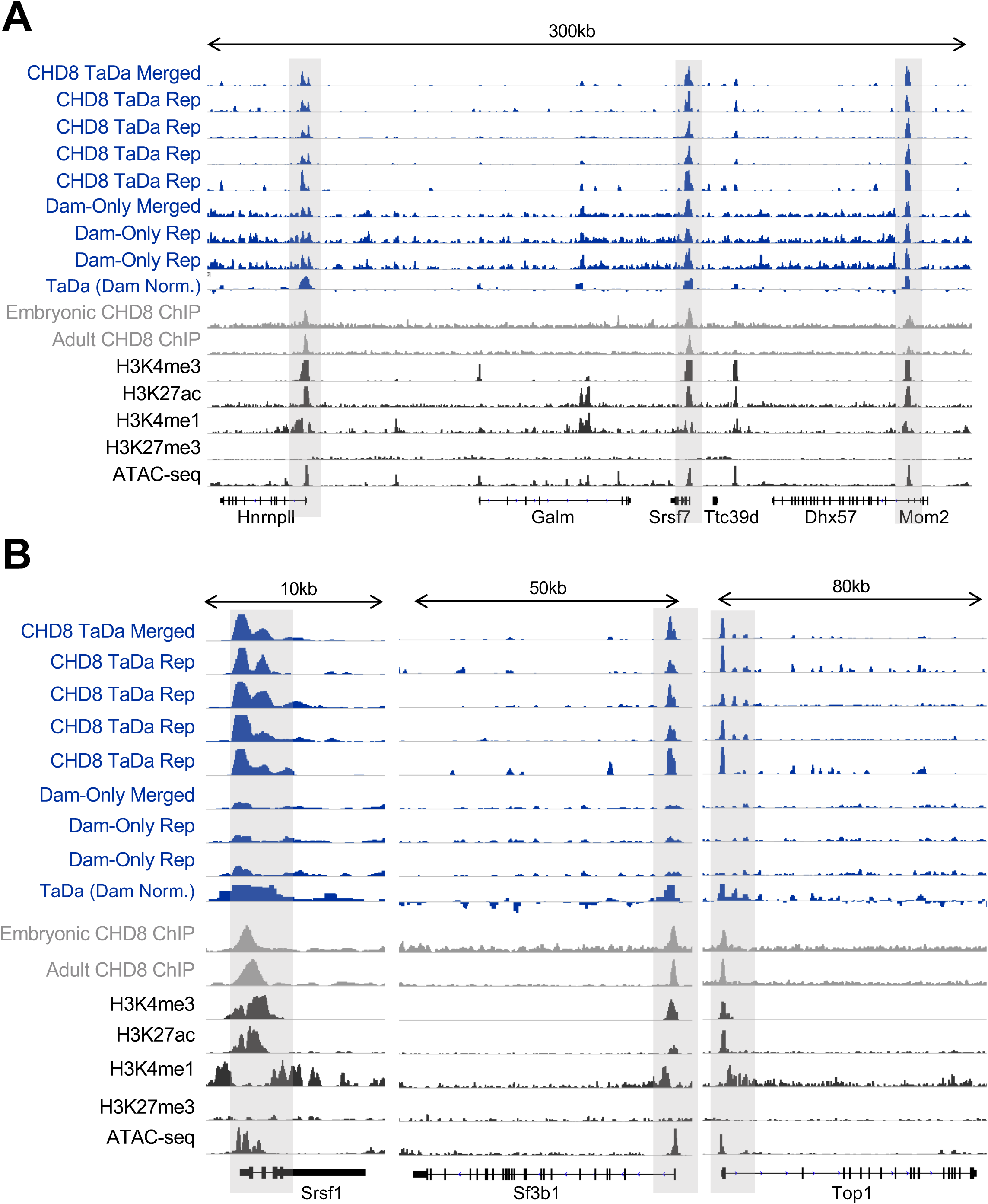
CHD8 binding near promoters across the genome across CHD8 TaDa-seq replicates. Loci featured in Figure 2 shown for all CHD8 TaDa-seq replicates. The first two CHD8 TaDa-seq replicates are the same replicates as in Figure 2. Data shows CHD8 binding at loci previously identified in CHD8 binding characterization studies, including RNA processing genes, *Hnrnpll* and *Srsf7* in panel A, *Srsf1* and *Sf3b1* in panel B, and a chromatin remodeling gene, *Top1*, in panel B. Grey boxes highlight CHD8 binding near identified promoters of interest. CHD8 TaDa-seq, Dam-only, or Dam-only normalized CHD8 TaDa (TaDa Dam Norm.) experiment tracks are in blue, CHD8 ChIP-seq experiments are in grey, and datasets of histone and chromatin accessibility signatures from the ENCODE consortium are in black. Linear representations of genes from the mouse mm10 genome are shown below coverage tracks. Height of the y-axis is scaled to show the peak for each track separately.

**Supplementary Figure S3.**
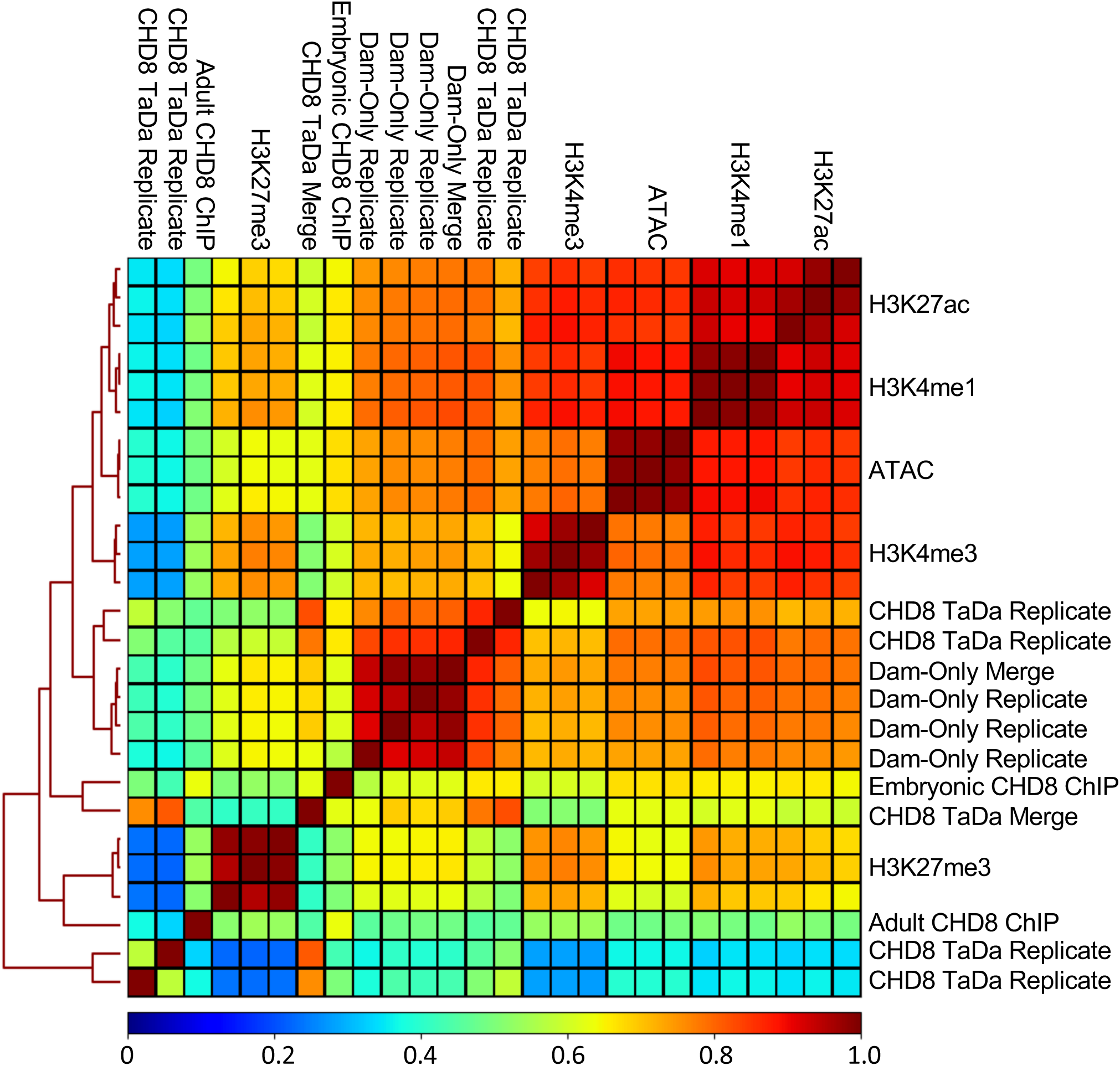
CHD8 binding is associated with open chromatin regions. Genome coverage correlation heatmap showing relationship between CHD8 TaDa-seq replicates, Chd8 ChIP-seq replicates, and ENCODE histone mark and chromatin accessibility dataset replicates. Data are hierarchically clustered according to similarity as indicated by a dendrogram. Differences in signal intensity between CHD8 TaDa-seq replicates in the correlation heatmaps were due to differences in sequencing depth. The legend indicates the correlation value between datasets. H3K27me3 is a histone mark associated with repressed DNA loci. H3K4me3 is a histone mark associated with actively transcribed promoters. ATAC-seq identifies regions of open chromatin. H3K4me1 and H3K27ac are histone marks associated with putative enhancers. CHD8 TaDa Merge – Merged CHD8 TaDa-seq dataset. Dam-only Merge – Merged Dam-only dataset.

**Supplementary Figure S4.**
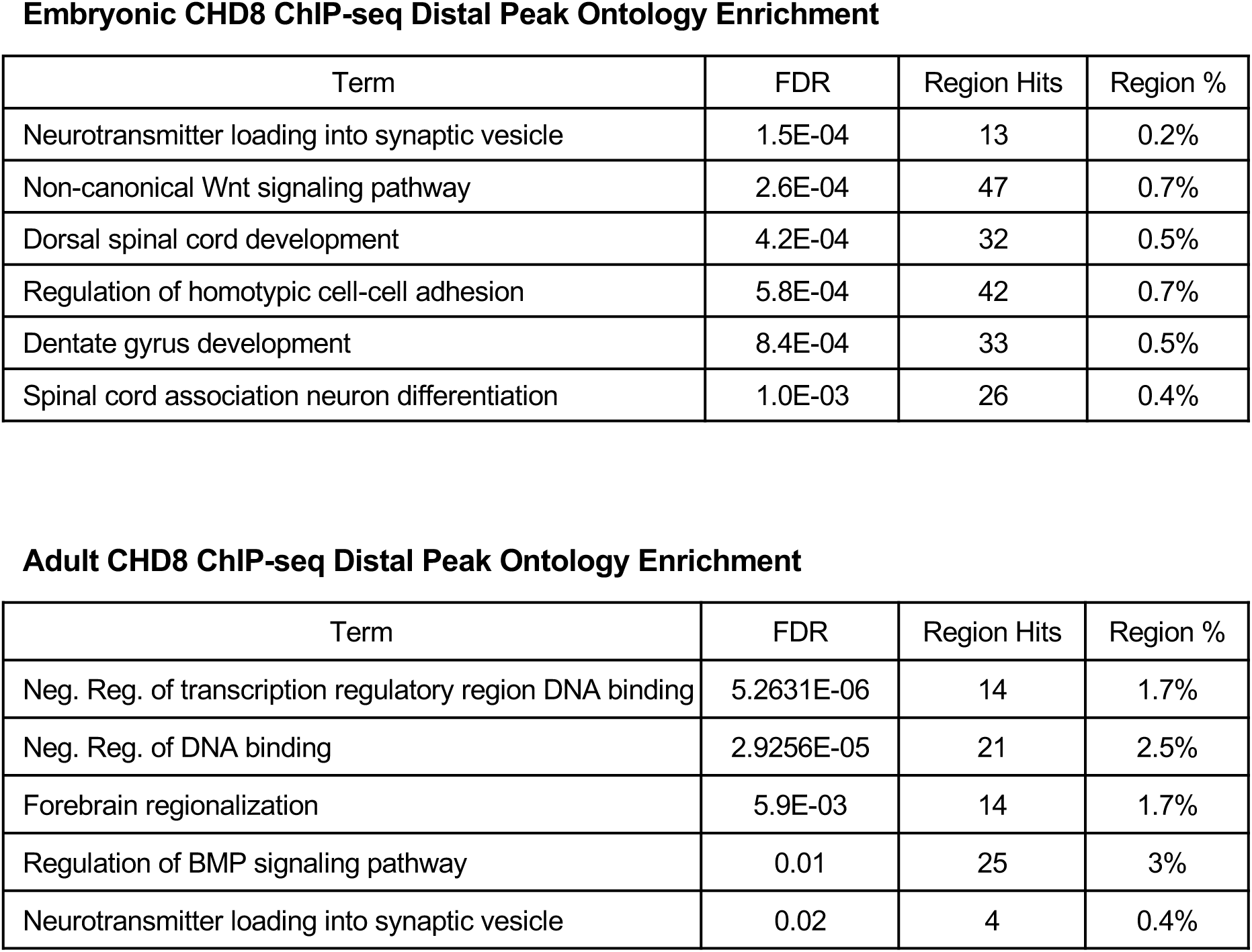
Evidence for promoter distal binding in CHD8 ChIP-seq datasets. Table showing functional annotations associated with peaks in CHD8 ChIP-seq datasets. Region hits capture the number of peaks associated with each term. Region % captures the percent of regions captured compared to the total number of peaks.

